# Morphologically Tunable Mycelium Chips for Physical Reservoir Computing

**DOI:** 10.1101/2025.08.20.671348

**Authors:** Orkan Telhan, Jake Winiski, Damen Schaak, Michael Siegel, Neale Petrillo, Eben Bayer

**Affiliations:** Ecovative LLC, Green Island, NY 12183

## Abstract

We introduce a neuromorphic computing substrate based on PEDOT:PSS-infused mycelium, a biofabricated, morphologically tunable material that can be engineered into electrically active components including resistors, capacitors, and non-linear elements. Leveraging the principles of physical reservoir computing, we demonstrate that hyphal networks grown under controlled environmental conditions can transform time-varying inputs into nonlinear, high-dimensional state trajectories, enabling machine learning tasks such as NARMA-10 sequence prediction. The chips are produced using a “design-grow-compute” workflow that integrates morphological modeling, parametric growth protocols, and vacuum-assisted polymer infusion. Morphological complexity is shown to influence charge transport and memory capacity, offering a new axis of control for designing analog computational architectures. Our prototype chips interface with a custom carrier board enabling analog signal conditioning and readout. Benchmarking revealed robust nonlinearity, temporal dynamics, and task-relevant separability. Unlike memristor arrays, photonic, or living-cell-based reservoir systems, our non-living analog mycelium chip is low-cost, biodegradable, and scalable using existing mushroom farming infrastructure, with production yields exceeding 3 million chips per growth cycle. This proof-of-concept demonstration using a single prototype device establishes the first biodegradable reservoir computing platform, with performance trade-offs justified by unprecedented sustainability advantages and orders-of-magnitude cost reductions. This advances a novel direction for biologically derived, single-use (compostable) or very large-scale machine learning hardware and introduces mycelium as a functional medium for analog inference.

## 1 Introduction

Physical reservoir computing is a framework that uses the noisy, transient, and non-linear aspects of physical systems to compute information [1, 2]. Its underlying premise is to use a network of dynamic reservoirs to process low-dimensional data into high dimensions and focus the training into a single readout layer. Historically, it is demonstrated that reservoir computing offers significant advantages in ML and AI applications [3, 4]. Unlike neural networks, in which the weights of each layer must be updated, reservoir computing requires only one layer to be trained, which eliminates both the computational cost for backpropagation and the need for GPUs for matrix manipulations. As the readout layer can be implemented in traditional CPUs, reservoirs must be efficient, versatile, low-power, and highly tunable. Reservoir systems can be trained with smaller datasets, fewer parameters and be suitable for online and continuous learning applications.

Physical reservoirs can range from photonic circuits to soft robotics, memristors, spintronic devices, which offer novel ways to process time-varying inputs into rich internal dynamics that encode temporal patterns, enabling tasks like signal classification, prediction, and control [5, 6, 7]. Such reservoirs promise a low-power, low-latency and continuous machine learning paradigm that can leverage the mechanical or thermodynamic properties of material systems [8]. Reservoir computers offer unconventional computing architectures that can be massively scaled or parallelized in different embodiments. The computational elements can be soft or rigid co-located or de-centralized [9].

While a variety of physical systems are evaluated as reservoir computers, not every physical non-linear system makes a good reservoir [10]. For example, a physical system should be able to give the same response when the same input is repeated [11]. The memory capacity (MC) of a reservoir also defines its ability to predict the same signal with different amounts of delay. Higher memory capacity indicates the system can retain and distinguish longer sequences of past input, making it better suited for time-dependent tasks like speech recognition, system identification, or time series forecasting.

Different physical reservoir designs which use biological, optical, electrical, magnetic, and mechanical components require different levels of manufacturing complexity and pose unique challenges for scalability. For example within the domain of Electrical RC, different memristor architectures provide non-linearity through modulating conductivity over analog resistive elements [12, 13, 14]. The memristor-based memory market has already reportedly reached $419 million in 2023, but challenges in achieving yield, reliability, device-level precision (high-level fixed point arithmetic) and high compute density currently slow down accelerating their use in-memory computing applications [15]. Memristor reservoir technology requires further development, and remains insufficiently mature for practical deployment as a replacement to conventional machine-learning infrastructure [16].

Mycelium, the vegetative structure of fungi, offers a novel and scalable medium for neuromorphic reservoir computing. Reservoir chips can be grown at existing mushroom farms without capital-intensive infrastructure or specialized tooling. In this paper, we demonstrate custom mycelium reservoir morphologies with tunable electrical properties—such as resistance, capacitance, and decay rate—designed pre-growth using a predictive model. We tested the feasibility of a model-driven “design-grow-compute” framework to grow mycelium chips using high-throughput benchtop reactors, which thermodynamically simulate the conditions in large scale farms.

The chips were grown in 19×14×4cm trays, cut to the desired dimensions, and vacuum-infused with PEDOT:PSS, a conductive polymer widely used in the thin-film industry.[17]. We have benchmarked different chip morphologies and evaluated their potential as reservoirs by assessing their separability, memory capacity, nonlinearity, and task accuracy. We quantified differences in morphological complexity and conductive dynamics as they transform incoming signals and conceptualize this transformation as a form of kernelization—a temporal, analog, nonlinear feature expansion. In this framework, a mycelium reservoir functions as a physical kernel, projecting input signals into a dynamic, high-dimensional feature space in which a simple readout can perform complex tasks such as classification [18].

When a physical kernel transforms an incoming signal via complex nonlinear dynamics, it implicitly maps that signal into a higher-dimensional space—without the need for symbolic or numerical encoding. In the context of mycelium reservoirs, the “kernel function” is physically instantiated through the organism’s morphology and conductive behavior. To design or tune such a kernel, we use a growth model that adapts the physical structure of the reservoir to suit a given computational task. Alternatively, the state of the kernel can be modulated by external stimuli such as temperature or electrical conductivity—though such dynamic tuning is beyond the scope of this paper.

Mycelium chips were fabricated with 16 spatially distinct reservoirs and mounted on a custom-designed carrier board that supports signal amplification, summation, and voltage level translation for analog interfacing (Figure 1). Each reservoir is electrically addressable, enabling parallel or sequential activation for diverse machine learning tasks. The modular architecture allows individual chips to be used independently or stacked to form larger, multi-reservoir computing arrays. While the intrinsic conductive properties of individual reservoirs are fixed post-fabrication, a form of functional learning can be implemented by dynamically routing input signals through subsets of reservoirs with predictable nonlinear responses. Using a multiplexing scheme, the system can reconfigure information flow toward nodes exhibiting desirable characteristics—such as sharp current transitions driven by space-charge-limited conduction (SCLC) [19, 20]. By exploiting these diode-like behaviors and known voltage-response boundaries, the network can be tuned to enhance task-specific performance without altering material weights, enabling adaptive computation through structural reconfiguration rather than parameter updates.

**Figure 1:**
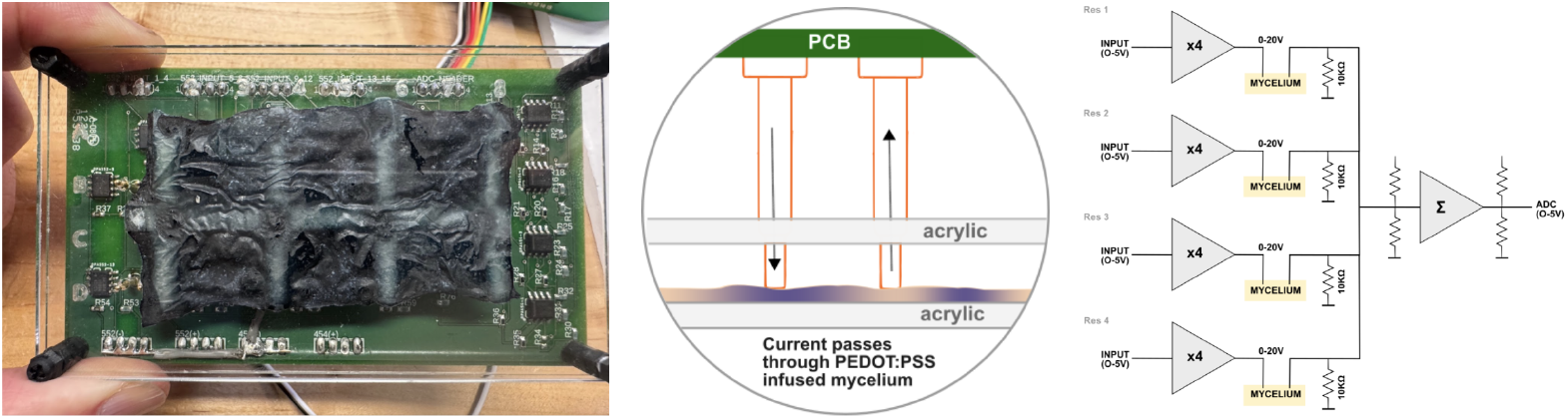
Neuromorphic Mycelium Chip mounted on a carrier board, which dispatches signals to 16 reservoirs. 4 sets of 4 reservoirs are summed with a summing amplifier and conditioned for ADC. A grid of pogo pins define locations of reservoirs, from which electrical signals are passed through mycelium.

To evaluate the computational capabilities of the mycelium-based reservoirs, we conducted a series of benchmark tests focused on temporal memory, nonlinearity, and sequence modeling. These included impulse response and correlation-based memory assessments, settling time characterization, and the NARMA-10 benchmark—an established nonlinear autoregressive task used to evaluate temporal information processing in physical reservoirs [6]. Mycelium reservoirs exhibited significant autocorrelation, cross-correlation, and nonlinear voltage responses consistent with memory-rich dynamics. When evaluated using regression models such as ridge regression and random forest, the reservoirs demonstrated predictive capability across delayed inputs, confirming their suitability for temporal inference tasks.

Our design required mycelium chips to be sandwiched between acrylic plates, and then mounted on a carrier PCB where digital or analog signals can be interfaced with the reservoir. This form factor allows multiple chips to be stacked and used for multiple machine learning tasks in parallel. In our experiments, the chips were not treated with any additional chemistry –shelf-stabilized using coatings or sealants. As an initial application area, we considered their applications in single-use or disposable electronics such as point-of-care diagnostics, food spoilage or hazard detection, and physical cryptography—where their tamper-proof construction and biodegradable composition ensure they are fully compostable within three months [21, 22, 23, 24]. Morphological tuning paired with post-growth infusion allows us to parametrically set the response intervals of the reservoirs, and even pave the path towards building multi-stage neural-network-like inference engines with set weights. In practical contexts such as packaging or environmental monitoring, these chips can be triggered by spoilage events (e.g., gas release or pH shift), classify contamination type and severity using embedded analog inference, and activate localized responses such as color change or signal dispatch. Mycelium chips can potentially be sealed against moisture loss or reinforced for durability with traditional tanning chemistry, enabling both extended field use and safe, compostable disposal.

Mycelium has been explored for its computational potential both as a living organism and as a material [25]. Adamatsky et al. demonstrated that live hyphal tissue can transmit signals through substrate and be used to build digital gates [26, 27]. Hyphal cells can also retain moisture and electrolytes in their network and store a reasonable amount of charge [28, 29, 30].

Flexible or rigid mycelium sheets also function as carrier surfaces to carry conductive traces and offer a biodegradable alternative to printed circuit boards (PCBs) [31]. Mycelium can be coated with copper using physical vapor deposition or electroplated with gold to achieve highly conductive traces (> 9 *×* 10^4^ S cm^*−*1^). Danninger also demonstrated that mycelium can be soaked with PEDOT:PSS to create vias that would allow multiple layers to be electrically bridged to create stackable circuits.

Filamentous fungi display remarkable phenotypic plasticity, dynamically remodeling their mycelial networks in response to spatial and environmental cues such as nutrient gradients and physical heterogeneity [32]. This plasticity is manifested through context-dependent modulation of branching, fusion, and growth direction, enabling decentralized adaptation for efficient resource acquisition, structural resilience, and morphological optimization. Morphological engineering in filamentous fungi in submerged culture systems involves the deliberate manipulation of growth form to enhance fermentation performance, leveraging the inherent phenotypic plasticity of these organisms [33].

Aerial mycelium refers to a cohesive, vegetative network of fungal hyphae that grows into open space above a nutritive substrate under solid-state conditions, forming a pure mycelial material without the development of fruiting bodies [34]. We morphologically engineered aerial mycelium by leveraging environmental strategies to control growth topology and tissue density. Specifically, we used topology adjustment layers—structured, perforated surfaces placed above the substrate—to regulate localized gas exchange and humidity, guiding hyphal emergence into geometrically defined patterns ranging from homogeneous to heterogeneous [35, 36]. The configuration of the growth chamber, including wall height and the application of casing layers, influenced internal gradients of CO_2_ and humidity, enabling directional control over vertical expansion and spatial uniformity [37]. Together, these methods enable scalable and repeatable engineering of mycelial structure.

## 2 Methods

### 2.1 Mycelium Production

Pure mycelium mats were produced according to Ecovative’s aerial mycelium platform using a benchtop bioreactor system, which under varying growth conditions, yielded a variety of intra- and inter-mat tissue morphologies. This platform leverages the natural plasticity of vegetative growth to engineer a wide morphological vocabulary, allowing for targeted variation in both organizational complexity and density within a single cohesive mycelial thallus. Within the scope of these studies, this provided a population of aerial mycelium mats from which various morphological classes of material could be selected. All mycelium mats were produced with Ecovative’s core foam strain and oven dried to *<* 10% moisture content prior to use, with stock panels up to 70mm in thickness (z-dimension) with arbitrary dimensionality in the x- and y-dimensions.

### 2.2 Mycelium Chip Specimen Preparation

Stock mycelium mats were sliced in the x-y plane using a commercial deli slicer to 1mm thickness, then further cut to 50×100mm in the x-y dimensions. The specimen was then misted with 70% isopropanol as a pre-wetting step and immediately floated in an excess of 1.3% stock solution of PEDOT:PSS (Ossila) in a reservoir plate (Corning HTS Transwell CLS4395). Specimens were then vacuum infused at -0.8 bar for 30 seconds (Vevor 5 gallon vacuum chamber with 3.5 CFM pump). After vacuum infusion specimens were removed from the reservoir plate and blotted with a paper towel to remove excess PEDOT:PSS.

Specimens were sandwiched between two sheets of acrylic and dried to 10% moisture content in a commercial dehydrator at 24 °C overnight. The ridges of the enclosure allowed PEDOT to be localized into a grid pattern as the pressure applied by the acrylic forced the polymer into non-pressurized areas.

### 2.3 Mycelium Chip Specimen Imaging and Morphological Featurization

All 50×100×1mm mycelium specimens were imaged under transmitted light using the film scanning function of an Epson V600 scanner, both before and after PEDOT:PSS infusion.

Morphological analysis was conducted via a custom Python pipeline using OpenCV and SciPy. High-resolution images were converted to grayscale, contrast-enhanced, and smoothed with Gaussian blur to improve feature clarity. Blob detection was performed using parameterized filters for area, circularity, convexity, and inertia, enabling quantification of individual morphological units. For each blob, diameter, area, and mean grayscale intensity were extracted, with summary statistics (mean, standard deviation, min, max, median) computed. Whole-image texture features were also derived from grayscale histograms, including contrast, energy, homogeneity, entropy, variance, skewness, kurtosis, autocorrelation, and difference entropy. This feature set captured both blob-level and global image characteristics for subsequent dimension reduction and correlation with electrical properties.

### 2.4 Mycelium Chip Electrical Characterization

For each specimen, resistance, capacitance, and voltage–current (I–V) curves were measured. Resistance was recorded in a matrix pattern using a Keithley 2400 source meter with 4-wire Keithley 5805 Kelvin probes at 2V, 16V, and 18V. Capacitance was measured at the same locations using an ET4410 LCR meter with a 1KHz sine wave (1V peak-to-peak, 100Ω input impedance). I–V curves were acquired using a Keithley 2400 with a full voltage sweep from 1-20V in 0.5V steps (Figure 2). All tests were repeated to ensure reproducibility, consistently revealing non-linear I–V behavior.

**Figure 2:**
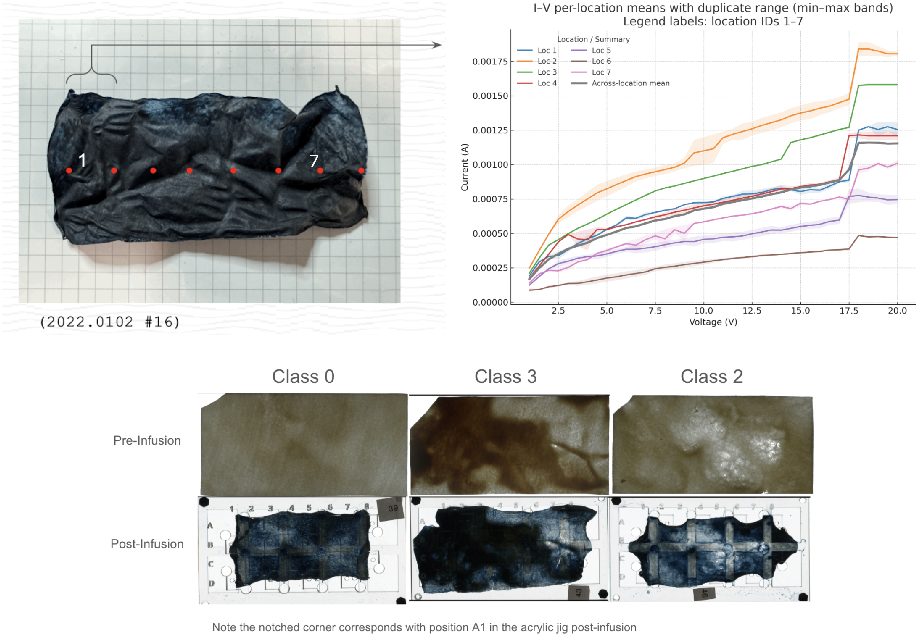
(Top) Current–voltage (I–V) characteristics across seven spatial locations with two technical sweeps per location. For each location, the band denotes the min–max across the two technical sweeps (n=2). The thick line overlays the across-location mean of per-location means (n=7). Error bands reflect technical-duplicate variation, not between-location variability. (Below) Representative chip specimens from clusters 0, 2, and 3, pre- and post-PEDOT:PSS infusion.

To calibrate each chip, electrical response was evaluated at eight locations by passing a 0–20V test signal through gold-plated prongs (3mm apart), arranged in a matrix, and recording voltage decay time.

### 2.5 Classification of Local Electrical Properties of Mycelium Chip Specimens as a Function of Global and Local Morphological Qualities

To identify distinct mycelium morphologies, aerial mycelium samples were processed through a morphological featurization pipeline, followed by PCA for dimensionality reduction and visualization, and k-means clustering for discrete group identification. Features were standardized using z-score normalization prior to clustering. The optimal number of clusters was determined via the elbow method (minimizing WCSS). From the resulting clusters, three were selected to maximize morphological diversity by choosing the pair with the greatest centroid distance and a third with the largest combined distance from the first two.

For controls, paper towel (Bounty Regular Roll, 2-ply) was soaked in PEDOT:PSS (Ossila PH1000; 10 mL bath) for 10 min and dried for 30 min at 24 °C. Rectangular specimens (2.54 × 2.54 cm) were cut and placed in isolated plastic wells. I–V curves were recorded in a four-wire Kelvin configuration (Keithley 2400), with probe tips centered and 3 mm apart; up sweeps only (1–20 V, 0.5 V step). Data are summarized as the median (min–max) across n = 5 replicates (independent pieces) at key voltages (5, 10, 15, 18, 20 V). The controls exhibited near-linear I–V behavior (median *R*^2^ = 0.995–0.999 over 2–16 V), whereas mycelium + PEDOT:PSS was less linear (*R*^2^ = 0.970–0.977) (See Supplementary Document Section 1.4)

Clusters 0, 2, and 3 were selected, and one representative specimen from each was prepared as chip specimens in acrylic templates for localized morphological and electrical evaluation.

Specimens were imaged before and after PEDOT:PSS infusion, cropped to final dimensions, and aligned using rigid body transformation in FIJI [38]. ROIs were defined from the post-infusion template and applied to the pre-infusion image to correlate electrical data with native morphology. Histogram-based image features (e.g., mean, skewness, entropy, contrast) were extracted per ROI, and whole-chip morphology was dimension reduced via PCA. Electrical data were processed by fitting linear models to IV curves and extracting slope, intercept, *R*^2^, and residuals. Skewed targets were transformed (Box-Cox/log) for model training (Figure 3).

**Figure 3:**
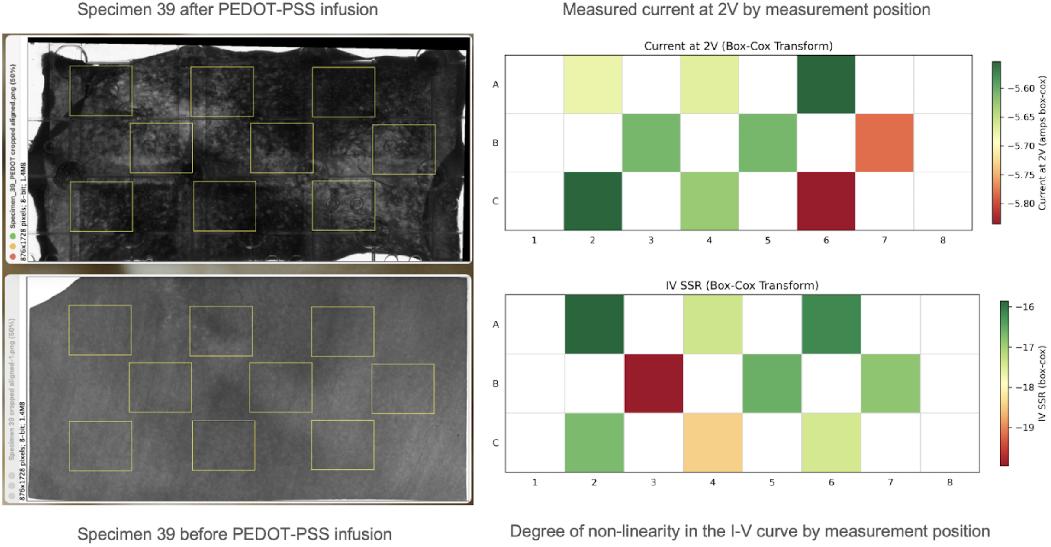
ROI alignment and mapped electrical features (class 0 specimen) showing spatial variation in current and I–V non-linearity.

PCA on electrical features revealed high covariability, with PC1 accounting for over 90% of variance. Mutual information analysis ranked predictive features, revealing strongest relationships with global morphology (density, complexity), followed by local density, texture, and eccentricity (Figure 4).

**Figure 4:**
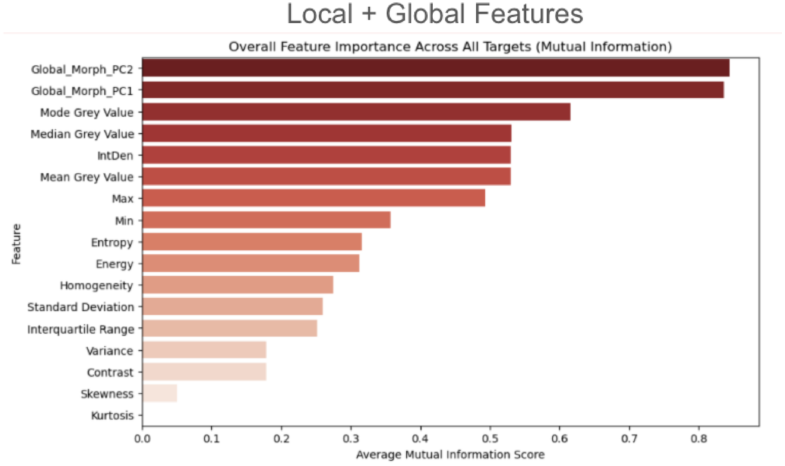
Mutual information highlights global and local morphological features of electrical behavior (PC1 and PC2 are global, all others are local

A feedforward neural network was trained to predict PC1 from global and local features using LOOCV. The architecture used two hidden layers (128, 64 ReLU units), L2 regularization, and dropout. The model achieved *R*^2^ = 0.74, indicating strong predictive ability (See supplemental information Fig. S14). To validate the systematic nature of morphology-property relationships, spatial autocorrelation analysis using Moran’s I confirmed significant organization of electrical resistance within the morphological feature space (p = 0.003-0.007 across test positions). SHAP analysis revealed morphological complexity and density (PC1 and PC2) as the strongest predictors of electrical resistance, exceeding the influence of PEDOT:PSS uptake (Supplemental Fig. S15). This hierarchy confirms that electrical properties are sensitive to morphological variation, and thus may be designed through morphological engineering independent of polymer characteristics, with structural complexity and density serving as the primary control parameters.

These results confirm that native mycelial morphology is highly predictive of post-infusion electrical behavior independent of the absolute mass uptake of PEDOT:PSS. This enables pre-infusion classification of electrical properties across a mycelium sheet, allowing for informed region selection to optimize reservoir computing function. By predicting electrical response from morphology, this approach supports rational design and reduced empirical trial-and-error in substrate engineering.

### 2.6 Design - Grow - Compute Workflow

Our approach enables a full-stack workflow for developing application-specific mycelium chips (ASMCs) that are designed in silico, biologically grown, post-processed, and directly interfaced with electronic systems (Figure 5). The process begins with feature selection, such as the number, spatial distribution, and electrical characteristics (e.g., resistance, capacitance) of reservoirs, guided by parameters derived from both global and local morphological feature spaces. These desired traits are translated into environmental growth parameters specific to the production setup (e.g., half-tray, full-tray, or bed-scale bioreactors). Mycelium tissue is typically grown in one- or two-week cycles, with multiple chip designs cultivated in parallel. After harvest, tissues are dried, precision-sliced, and imaged. These pre-infusion images are analyzed to extract morphological features and feed back into predictive models for iterative design refinement. Sliced specimens are mounted into acrylic cartridges and infused with PEDOT:PSS, as described in the Methods section; each cartridge is referred to as a single chip. Chips are then mounted onto carrier boards for electrical characterization. A calibration routine pulses each reservoir at various voltage levels to baseline its dynamic response characteristics, preparing the chip for deployment in analog computing tasks.

**Figure 5:**
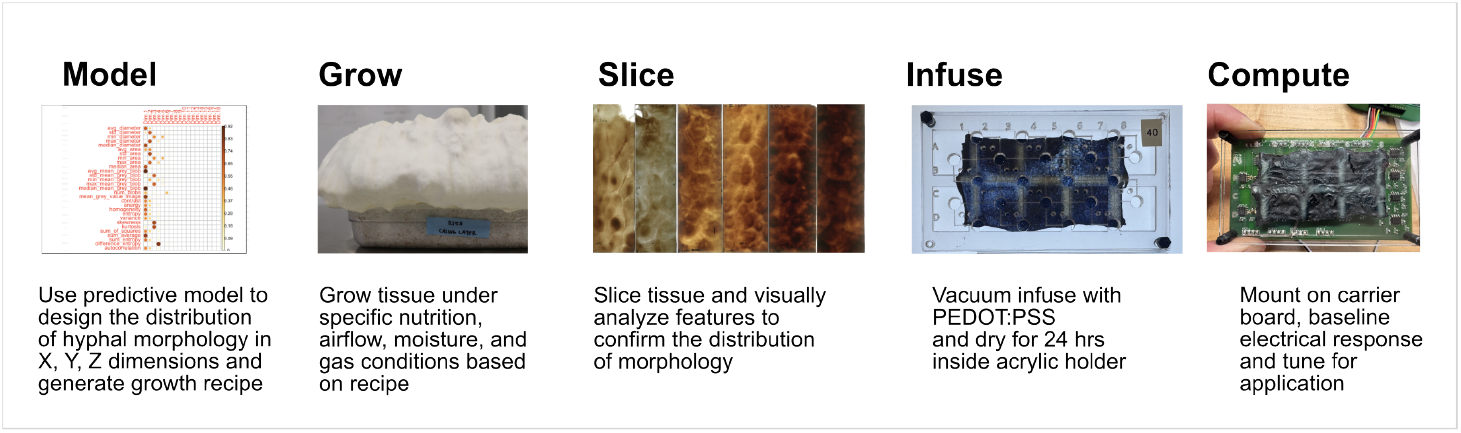
Full-stack workflow for producing application-specific mycelium chips (ASMCs), from in silico design through modeling, growth, slicing, infusion, electrical calibration, and computation steps.

## 3 Results

### 3.1 Computing with Neuromorphic Mycelium

We designed the carrier board that serves as an analog interface to the reservoir network, routing both input signals and their corresponding collection pathways. Sixteen independent input channels operate in parallel, enabling the configuration of reservoirs independently [39]]. Additionally, multiple channels can be used sequentially to define reservoir sub-clusters composed of several inputs. The outputs from four channels are summed via a non-inverting operational amplifier, which also conditions the signals for acquisition by an analog-to-digital converter (ADC). Multiple carrier boards can be networked together to construct more complex reservoir layouts.

The input to the mycelium chip can be digital–from the output of a microcontroller, or analog–a waveform/signal directly sourced from a sensor. For our prototypes we used Digilent’s Analog Discovery 2 and 3 devices with both continuous and discrete input to test the capabilities of the reservoirs. The carrier board offers a two stage amplification process. Each signal from 0-5V range is amplified with a non-inverting amplifier with 4x gain to achieve the critical 16-18V threshold that causes the non-linear jump in current flow. After the summation phase, the signals are stepped down with a voltage divider back to 0-5V range and buffered for the ADC.

We have evaluated the chip’s computational capabilities on two fronts: memory response and temporal processing capacity. Our experimental setup used the analog carrier chip that received input from one Analog Discovery 2 (channels 1&2) and one Analog Discovery 3 Device (channels 1). The summed output of three channels were read into the Analog Discovery 3 device. We used 3 reservoirs–locations A1, A4, and A6 on 3 chips on the same task.

### 3.2 Memory Response

We assessed mycelium reservoir’s temporal dynamics by applying three input types—random pulses, step signals, and sine waves—with programmable delays (Table 1). These patterns were selected to probe distinct memory regimes across three reservoirs in one chip.

**Table 1:** Temporal dynamics across input types for a single device, summarizing memory and response characteristics. “Max Autocorr (lag)” is the peak Pearson autocorrelation of the state at positive lags; “Max Crosscorr (lag)” is the peak Pearson correlation only; *R* between input_*t−*lag_ and state_*t*_ over non-negative lags. Lags are in seconds from the median sampling interval.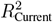 uses input at 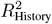 uses inputs *t*, …, *t −* 10. “—” indicates not observed within the recorded window.

| Input Type | Max Autocorr (lag) | Max Crosscorr (lag) | $R^2$ (Current) | $R^2$ (History) | $\Delta R^2$ | Settling ($\pm 10\%$) [s] |
| --- | --- | --- | --- | --- | --- | --- |
| Step (K=2 runs) | 0.982 (3.15 s) | 0.472 (11.10 s) | 0.2556 (95% CI 0.0000–0.5113) | 0.3705 (95% CI 0.0000–0.7411) | 0.1149 (95% CI 0.0000–0.2298) | — |
| Random (K=9 runs) | 0.342 (2.50 s) | 0.721 (0.00 s) | 0.185 (95% CI 0.085–0.283) | 0.292 (95% CI 0.140–0.437) | 0.107 (95% CI 0.048–0.166) | — |
| Sine (K=3 runs) | 0.497 (1.00 s) | 0.362 (1.00 s) | 0.1844 (95% CI 0.0000–0.4698) | 0.2296 (95% CI 0.0000–0.4748) | 0.0452 (95% CI 0.0000–0.1305) | — |

For random inputs, adding short input history improved next-step state prediction within-device (mean Δ*R*^2^ = 0.107, *p* = 0.010), consistent with reliable carryover on sub-second to 1 s scales. The state showed modest persistence (median peak autocorrelation = 0.342 at 2.5 s) while maintaining strong input coupling (median cross-correlation = 0.721 at 0 s). Under sinusoidal drive, the state exhibited phase-locked memory aligned to the waveform. Two of three runs showed clear phase-locked input–state coupling; one run was non-responsive, reflecting device variability. Step inputs revealed multi-timescale dynamics with both rapid initial response (*τ ≈* 0.15 s) and slower drift components (10–90% rise *≈* 4.35 s) that prevented settling within recorded windows

The device thus expresses a spectrum of memory behaviors: transient under stochastic drive, phase-locked under periodic drive, and fast-plus-slow following steps. This responsiveness supports suitability for time-dependent analog computation. However, all findings represent technical replicates on one chip (biological *n* = 1; establishing generality will require replication on independent devices. See Supplemental Section 1.5 for details and definitions.

### 3.3 NARMA-10 Benchmark

To evaluate the reservoir’s temporal processing capacity, we employed the standard Nonlinear Auto-Regressive Moving Average task of order 10 (NARMA-10)—a widely used benchmark for assessing nonlinear memory in physical reservoir systems [6, 40]. This task requires the reservoir to predict the next value in a nonlinear time series based on the preceding 10 inputs, offering a stringent test of both memory depth and nonlinear transformation.

We used the same chip and three reservoirs for testing, generating 1,000 training samples by applying voltage signals in the 0–5 V range. Feature representations included the raw reservoir state, its square, and trigonometric expansions (sin(state *×* 3), cos(state *×* 2)). These enriched features were used to train both Ridge regression and Random Forest models across multiple reservoir configurations.

Our mycelium reservoir system achieved NARMA-10 performance across three independent test experiments with Ridge regression yielding RMSE values of 0.102–0.106, corresponding to normalized root mean squared error (NRMSE) of 1.01–1.07, where NRMSE = RMSE*/σ*(target). Additional technical replicates on a single device (10 runs) showed mean NRMSE = 0.984 (95% CI 0.943–1.032) with mean *R*^2^ *≈* 0.027, indicating that the reservoir state carries short-term history useful for NARMA-10, though absolute accuracy remains near the mean-predictor baseline.

While this performance falls below optimized electronic reservoir systems (NRMSE *∼* 0.056–0.410), it represents the first demonstration of temporal computing in a biodegradable, agriculturally scalable substrate. The NARMA-10 evaluation revealed consistent performance reproducibility, successful nonlinear feature extraction from mycelial network dynamics, and stable operation within the 16–18 V nonlinear regime.

The demonstrated NARMA-10 capability validates the core concept of mycelium-based neuromorphic computing and provides a foundation for morphological optimization strategies. This performance is achieved at *<* $1 per chip with complete biodegradability—representing orders of magnitude cost savings for a bio-based reservoir computing platform. The consistent performance across different readout models supports the viability of mycelium reservoirs for applications where sustainability, cost, and biological integration are prioritized over absolute computational performance. This work establishes a new paradigm for environmentally sustainable neuromorphic computing with clear pathways for performance enhancement through morphological engineering.

## 4 Discussion

### 4.1 Operational Range and Durability of Mycelium Chips

While conventional AI chips operate at sub-1V core voltages to minimize power consumption and thermal load [41], our analog mycelium-based reservoirs require signal amplification to 16–18V to activate non-linear conduction regimes. This elevated voltage reflects the intrinsic charge transport behavior of PEDOT:PSS within porous mycelial matrices, where trap-state saturation consistent with space-charge-limited conduction (SCLC) enables diode-like transitions essential for analog inference [42]. Though uncommon in CMOS-scale devices, such operational ranges are acceptable in low-power, passive systems, especially when intended for single-use or intermittent-use applications where energy consumption is limited and bio-compatibility, biodegradability, and cost are the dominant design constraints. The platform is also compatible with energy scavenging or capacitor-driven systems under field conditions.

Mycelium chips were evaluated in ambient conditions (22°C unpowered; 36°C during operation, due to carrier board heating). Across multiple cycles, voltage response curves showed modest drift, likely due to moisture exchange and morphological reconfiguration. No sealing or moisture-stabilizing treatment was applied to these prototypes, so gradual conductivity decline over time is expected. However, baseline recalibration, range re-mapping, and voltage correction factors allow compensation for mild aging effects. In over 50 tests per chip, performance remained stable and repeatable, suggesting practical durability within the intended use window.

### 4.2 Scalability Potential

Our mycelium reservoir system achieves functional NARMA-10 performance (NRMSE = 0.984) and shows that biodegradable mycelium substrates can support temporal computing: the device tracks structured inputs (sine), retains short history under stochastic drive, and solves a standard nonlinear benchmark (NARMA-10) with simple readouts. While accuracy lags optimized electronic reservoirs, the platform offers compelling advantages in sustainability and cost (materials <$1 per chip) —a 500-50,000x cost advantage over competing technologies ($500-50,000+)—while uniquely offering complete biodegradability and agricultural-scale manufacturing.

Compared to other neuromorphic platforms, mycelium-based reservoirs offer a unique combination of biological integration, morphological tunability, and ultra-low fabrication cost. While memristor arrays provide superior performance with non-volatile dynamics at higher speeds, they require precision fabrication and face scaling challenges (Table 2). Photonic reservoirs offer excellent computational performance (NMSE 0.014-0.127) and unmatched bandwidth but remain difficult to fabricate and scale beyond lab prototypes.

**Table 2:** Performance and cost comparison of physical reservoir computing systems. Reported NRMSE values may differ by normalization method. Missing entries indicate unavailable or incomparable data. Cost and power estimates reflect material, fabrication, and operational differences across architectures.

| System | Power Consumption | NARMA-10 NRMSE | Processing Speed | Scalability | Sustainability | Cost Estimate |
| --- | --- | --- | --- | --- | --- | --- |
| **Mycelium Reservoir** | 2.5–30mW | 0.984 (*this work, mean RMSE across 10 runs) | 0.1–1s settling | Agricultural scale | Biodegradable, compostable | <1/chip, ~50/carrier |
| **Memristor Arrays** | 0.005mW@MHz<br>5nJ–4.28aJ [48] | 0.00361 [4] – 0.036 [49] | MHz–GHz | Fab-limited | E-waste concern | \$500–5,000+ |
| **Photonic RC** | 5100mW [50] | 0.0143 [50], 0.127 [51] | 60GHz | Lab/specialty fab | Traditional materials | \$5,000–50,000+ |
| **Ion-Gating RC** | Ultra-low power | 0.00938 [52] | Not reported | Nanodevice scale | Traditional materials | \$1,000–10,000+ |
| **Spin Wave RC** | 14.2W@50Hz [53]<br>284μJ/input | 0.168 [53] | 50MHz (20ns response) | Research scale | Traditional materials | Non-commercial |

These findings establish a practical starting point for bio-based reservoir computing, with clear levers for improvement—morphology and electrode design, richer feature/readout sets, and input protocols. Because the present evidence is within-device, the next step is replication across independent chips with pre-specified QC and standardized metrics; we expect this will sharpen performance estimates and accelerate optimization while preserving the system’s environmental benefits.

Unlike previous bio-inspired or living-biology reservoir computing approaches [43, 44, 45, 46], our work explores non-living mycelium as a highly scalable substrate. We demonstrated that electrical properties reflect morphological variation across chips *R*^2^ = 0.74, with morphological development controllable through biomanufacturing parameters. This suggests that electrical properties influencing reservoir characteristics may be designed into mycelium chips during the biofabrication process independent of the inherent characteristics of PEDOT:PSS by targeted morphological engineering. Specific tissue density and organizational characteristics can be programmed by modifying environmental conditions during growth (Figure 8).

**Figure 6:**
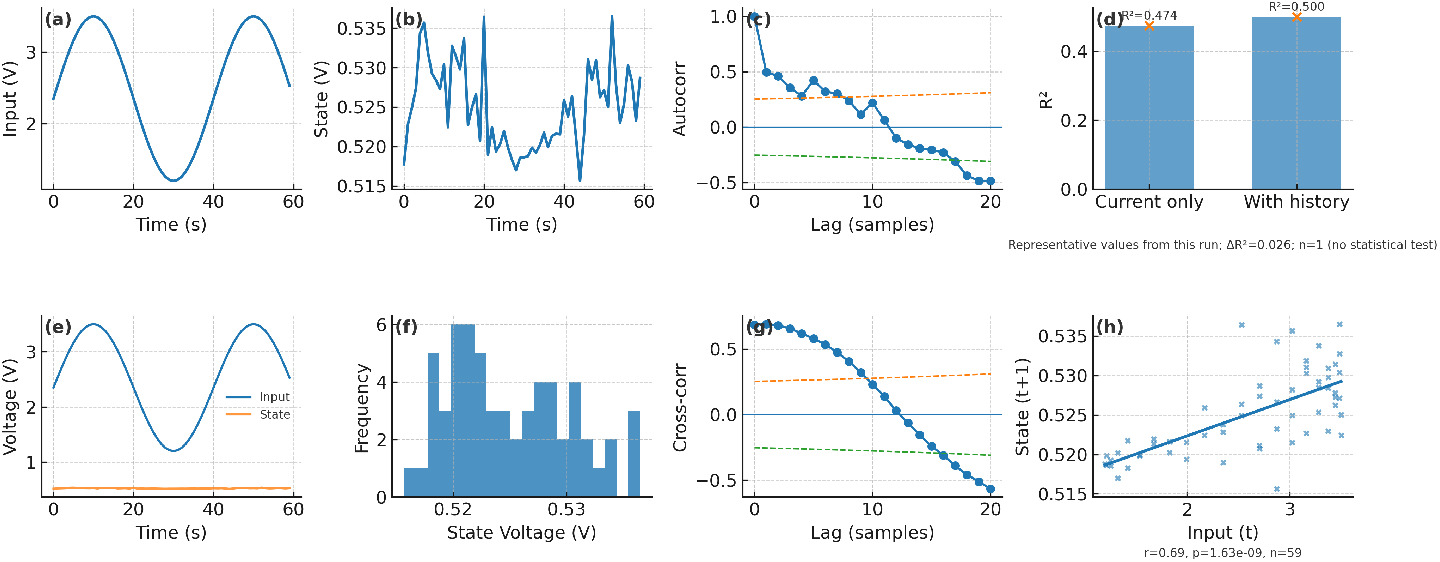
Temporal memory in a mycelium reservoir with sine input (representative device, biological *n* = 1). (a) Input waveform (mean; shaded 95% CI). (b) Reservoir state (mean; shaded 95% CI). (c) State autocorrelation (unbiased) with per-lag white-noise significance bounds 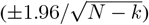. (d) Prediction performance (*R*^2^) for models using current input only vs. input with history from the same run: 0.4745 vs. 0.5005 (Δ*R*^2^ = 0.026). *n* = 1 (representative), so no inferential statistics are reported. (e) Representative time series from this device (one run). (f) Distribution of reservoir state values (all timepoints in this run). (g) Input–state cross-correlation for non-negative lags (0 … *K*) with per-lag white-noise bounds 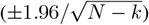. (h) Input at time *t* vs. state at time *t* + 1 with ordinary least-squares fit; panel annotates Pearson’s *r*, exact *p*, and *n* for this run.

**Figure 7:**
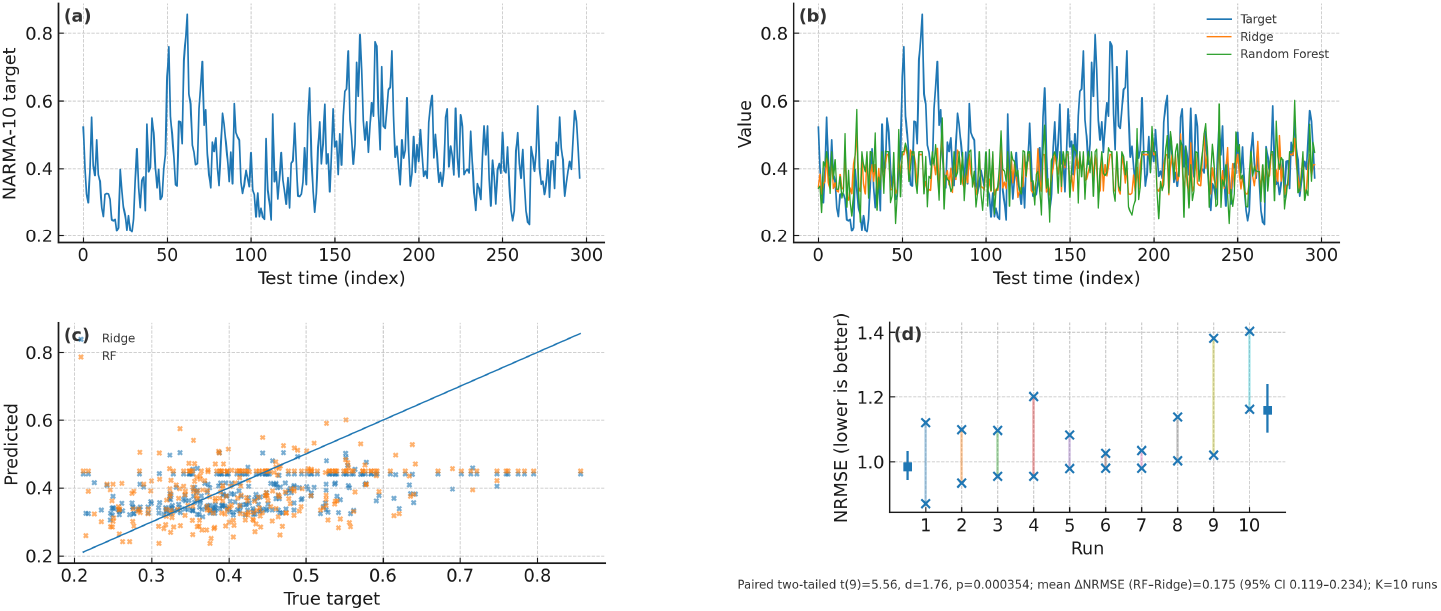
NARMA-10 evaluation on a mycelium reservoir (single device; 10 runs). (a) Target sequence segment from a representative run (held-out test). (b) Target vs. predictions from Ridge and Random Forest on the same run. (c) Predicted vs. true scatter for the representative run (45^*°*^ line shown). (d) Across-run comparison: paired dots show NRMSE for Ridge vs. Random Forest per run (lower is better); squares indicate mean *±* 95% CI across runs. Paired two-tailed *t*(9) = 5.56, *d* = 1.76, *p* = 0.000354; mean ΔNRMSE (RF–Ridge) = 0.175 (95% CI 0.119– 0.234); *K* = 10 runs.

**Figure 8:**
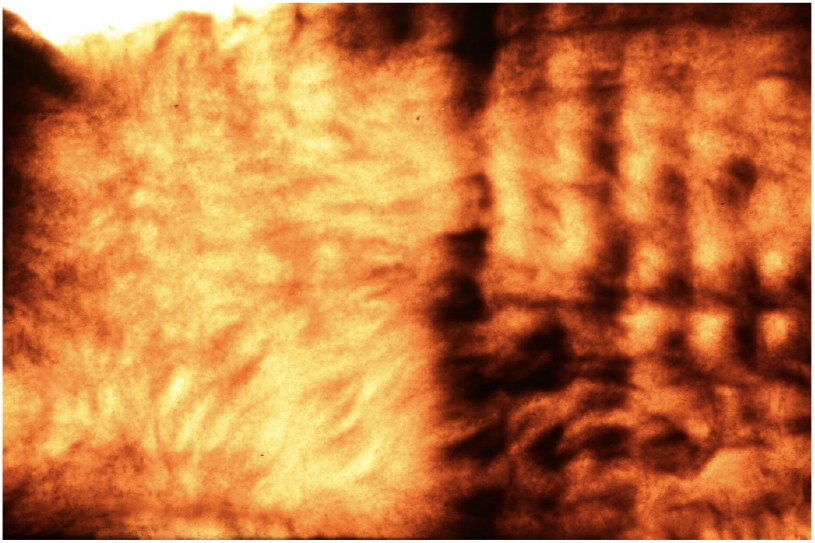
Various mycelium chip morphologies are mycofabricated within a single contiguous 8×12 inch sheet using aerial mycelium morphological engineering techniques (shown by transmitted light photography after slicing).

This work introduces a novel biomanufacturing paradigm for the first disposable ML inference chip. Aerial mycelium here functions simultaneously as substrate, signal processor, and morphological kernel. Analogous to panelized PCB manufacturing, a single aerial mycelium panel can embed diverse, task-specific morphologies in discrete regions. A standard Dutch-style mushroom bed (1.34 × 40 m) can produce over 3 million chips per 7-14 day cycle [47], enabling a fully integrated, biodegradable hardware stack for sustainable, task-specific inference where cost and environmental impact outweigh absolute computational performance. Building on recent demonstrations of conductive mycelium circuit substrates [31], we envision a future in which aerial mycelium serves as both substrate and computational medium, enabling a fully integrated, biodegradable hardware stack for sustainable, task-specific inference.

## Acknowledgments

This research was developed with funding from the Defense Advanced Research Projects Agency (DARPA). The views, opinions, and/or findings expressed are those of the author(s) and should not be interpreted as representing the official views or policies of the Department of Defense or the U.S. Government.

## Data Availability

The datasets generated and analyzed during the current study are available from the corresponding author upon reasonable request.

## Code Availability

All code used for signal processing and reservoir evaluation is available at https://github.com/otelhan/NeuromorphicMyceliumChip

## Supplementary Information

See Appendix below for additional figures and method discussion.

