## Supplemental Information for "Morphologically Tunable Mycelium Chips for Physical Reservoir Computing"

### 1 Supplementary Information

#### 1.1 Mycelium Chip Specimen Imaging and Morphological Featurization

All 50x100x1mm mycelium specimens were imaged via transmitted light using the film scanning function of an Epson V600 scanner, with image capture performed both before and after PEDOT:PSS infusion. Morphological analysis of transmitted light images was performed using a custom image processing pipeline implemented in Python, with OpenCV [1], and SciPy [2]. High-resolution images of each specimen were converted to grayscale, contrast-enhanced, and smoothed using Gaussian blur to optimize feature visibility. Blob detection was applied using parameterized filtering on area, circularity, convexity, and inertia, enabling quantification of individual morphological units. For each blob, diameter, area, and mean grayscale intensity were extracted, and summary statistics (mean, standard deviation, minimum, maximum, and median) were calculated. Additionally, whole-image texture features were computed from the grayscale histogram, including contrast, energy, homogeneity, entropy, variance, skewness, kurtosis, autocorrelation, and difference entropy. The resulting feature set captured both local (blob-level) and global (image-level) structural characteristics for downstream dimension reduction and morphological-electrical property analyses.

#### 1.2 Mycelium Chip Electrical Characterization

For each specimen we measured resistance, capacitance and voltage-current (V–I) curves. Resistance across the specimen was measured in a matrix pattern Keithley 2400 source meter using 4-wire Keithley 5805 Kelvin probes at 2V, 16V and 18V reference voltages. Similarly, capacitance was measured using the ET4410 LCR meter also at the same locations with a sign wave signal (Frequency @ 1kHz, Input Impedance @ 100 Ohms, Voltage @ 1V peak-to-peak.Voltage-current (IV) response was measured using Keithley 2400 source meter with a full voltage sweep (1–20V, with .5V step increments) (Figure S1). Tests were repeated twice for each specimen to validate findings. We consistently found a non-linear relationship in all specimens.

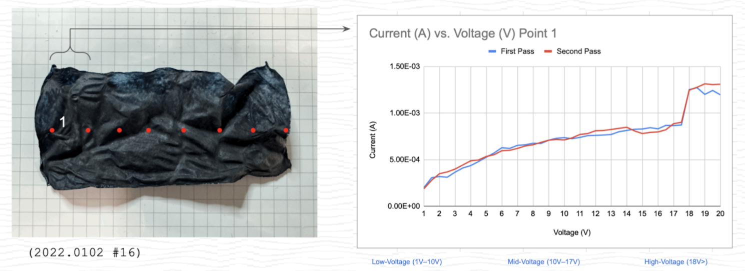


Figure S1: Electrical characterization of mycelium specimens shows consistent non-linear I–V behavior across all samples, measured via resistance, capacitance, and voltage sweeps.

Morphology influences PEDOT absorption and causes non-linearity in conductivity. We leverage the electrical differences exhibited at the different parts of the chip to model an electrical reservoir network for computation. We defined reservoirs as electrically addressable areas, or regions of interest (ROI), on the mycelium chip in which current can be passed and read through two pins. As electricity goes through the hyphal network there is a non-linear voltage drop across the pins, which can be attributed to the variable

We observed that hyphal structures exhibited a non-linear (diode-like) behavior between 16-18V on all tested locations on the surface. Mapping all different locations' response rate allowed us to map a dynamic reservoir network that can exhibit a delay (memory effect). We attributed this critical transition point as space-charge limited conduction (SCLC) in which the traps within the hyphal networks fill in and cause increased current flow.

To be able to commission a chip, we ran a calibration process to evaluate the electrical response of mycelium at 8 locations. We passed a 0-20V test signal through gold plated prongs (spaced 3mm apart) arranged in a matrix to measure the voltage and its decay time.

#### 1.3 Global Electrical Resistance of Mycelium Chips as a Combined Function of PEDOT:PSS Uptake and Mycelium Morphological Qualities

Mycelium chip specimens were selected to represent a cross-section of morphological variance reflecting a subset of the total 'morphological vocabulary' that is addressable through tuning aerial mycelium growth parameters. Figure S2 shows the transmitted light image of each included specimen annotated with its respective characterized electrical resistance after PEDOT:PSS infusion.

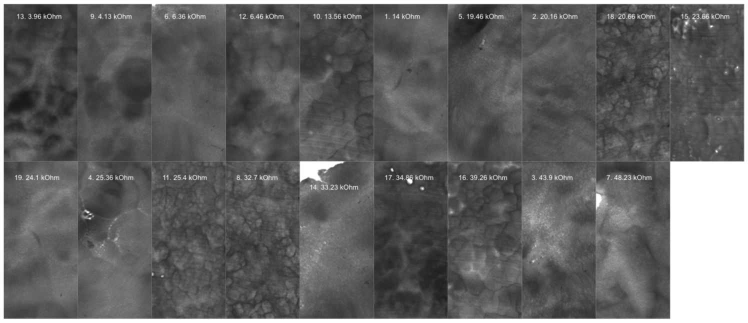


Figure S2: PEDOT:PSS infused mycelium morphologies characterized with lightbox imaging and resistive measurements.

First, the relationship between absolute uptake mass of PEDOT:PSS and electrical resistance was evaluated. It was found that there was a significant (p=0.5–0.004) negative linear relationship between absolute uptake mass of PEDOT:PSS and resistance at all test positions, but with low correlation ($R^2 = 0.2$–$0.41$). Fig S3 shows the linear correlation between absolute PEDOT:PSS uptake mass and measured resistance at each of four test positions.

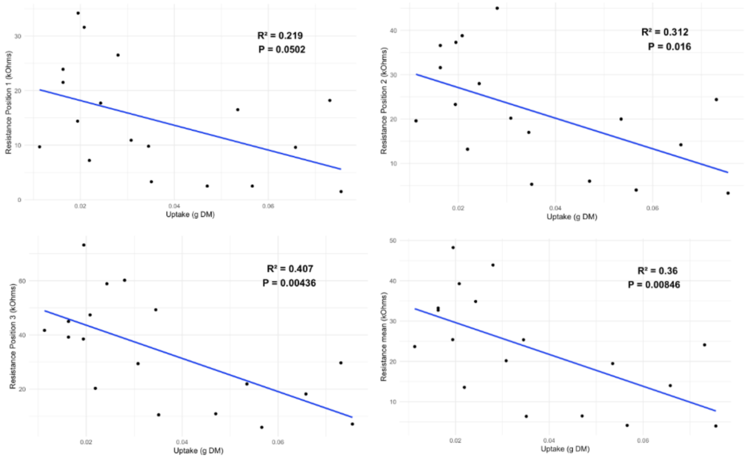


Figure S3: Correlation between absolute PEDO:PSS uptake and resistance.

This observed negative linear relationship between electrical resistance and absolute mass uptake of PEDOT:PSS is consistent with the role of PEDOT:PSS as a conductive dopant that enhances charge transport within the mycelium matrix. As the amount of infused PEDOT:PSS increases, a greater number of conductive pathways are formed, reducing the overall resistivity of the composite material.

Further evaluation of the relationship between mycelium mat morphology and residual resistance not explained by PEDOT:PSS mass uptake was performed (Figure S4). Dimension reduction of the resultant total morphological feature space was performed via principal component analysis, then spatial autocorrelation of residual resistance within the k-dimensional morphological space was used to evaluate the degree of organization of electrical resistance according to morphology.

Our cluster shows residual resistance (at measurement position 1) within the k-dimensional morphological space defined by principal components 1, 2, and 3 (representing >70% of total morphological variance). In this case a Moran's I p-value of .003 provides evidence of significant organization of residual resistance variance within the morphological feature space [3, 4]. Further testing of residual resistance at test positions 2 and 3 also supports evidence of significant autocorrelation within the morphological feature space (p=.007 and .005, respectively).

To further evaluate the relative importance of morphology for explaining resistance independent of tissue density and PEDOT:PSS uptake, feature importance was performed by:

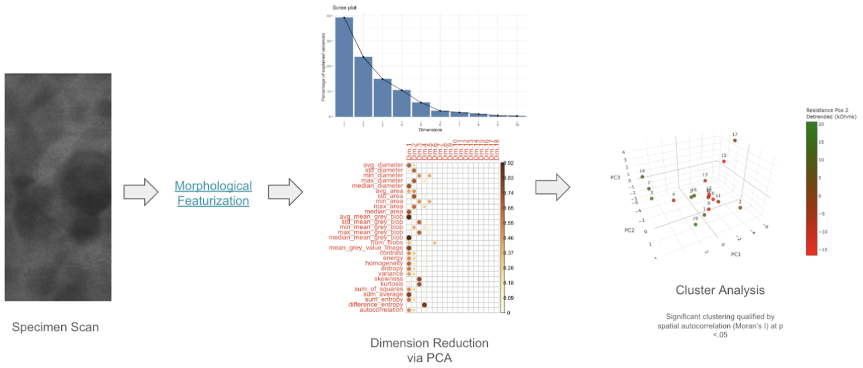


Figure S4: Specimen morphology analysis workflow.

1. Training a regression neural network (TensorFlow)[5, 6, 7, 8] that attempts to explain mean resistance of all four test positions as a combined function of pre-infusion specimen density, the absolute mass uptake of PEDOT:PSS, the uptake rate of PEDOT:PSS (g uptake per g tissue mass), and morphology (represented by principal components 1-4, capturing >80% of total morphological variance).
2. Reasonable predictive quality was resolved (coefficient-of-determination = 0.54), so feature importance was performed via SHAP (SHapley Additive exPlanations)[9] to evaluate the relative importance of morphology for explaining resistance, then,
3. LIME (Local Interpretable Model-agnostic Explanations) was used to explore what characteristics best explain the lowest and highest resistance specimens in the sample set [10].

The plot in Figure S5 shows the results of SHAP analysis showing the relative importance of each feature for explaining mean resistance (from top to bottom). Morphological PC1 demonstrated the strongest explanatory power for mean resistance followed by absolute mass uptake of PEDOT:PSS and morphological PC3 and PC4. This would suggest morphology is strongly related to resistance in combination with PEDOT mass uptake. Lower explanatory power is also associated with tissue density independent of morphology and the uptake rate of PEDOT, followed by morphological PC2. Generally, this suggests morphology is potentially impactful on resistance performance independent of (1) mass uptake of PEDOT:PSS and (2) tissue density. Further, there is sensitivity to multiple independent dimensions of morphology, potentially suggesting more complex relationships.

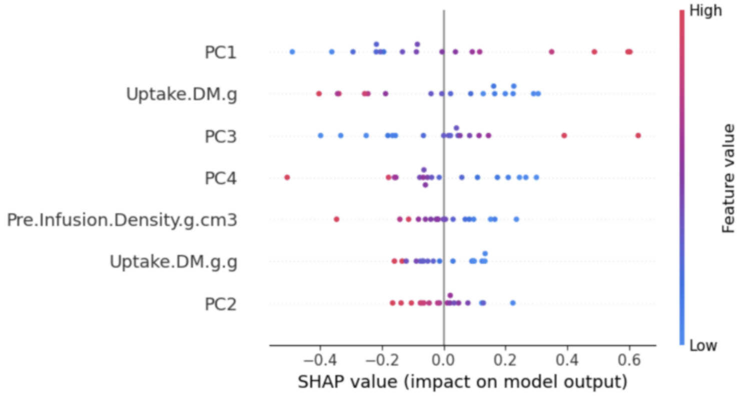


Figure S5: SHAP analysis reveals that morphological complexity, particularly PC1, is the strongest predictor of mycelium chip resistance, exceeding the influence of PEDOT:PSS uptake and tissue density.

Furthermore, the results of LIME analysis suggest which features are most critical for explaining the lowest (specimen 13) and highest (specimen 7) resistance (Figure S6). The results suggest attaining a low resistance is dependent on first attaining a high PEDOT:PSS uptake mass, then a particular morphology, followed by a high tissue density. Alternatively, attaining a high resistance is first dependent on morphology then a reduced PEDOT:PSS uptake mass.

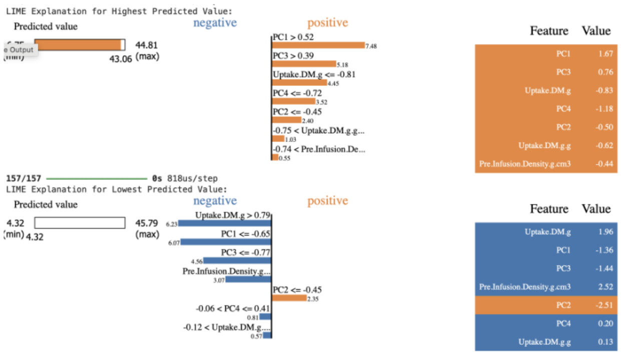


Figure S6: LIME analysis identifies PEDOT:PSS uptake as the primary driver of low resistance and morphology as the dominant factor in high resistance across mycelium chip specimens.

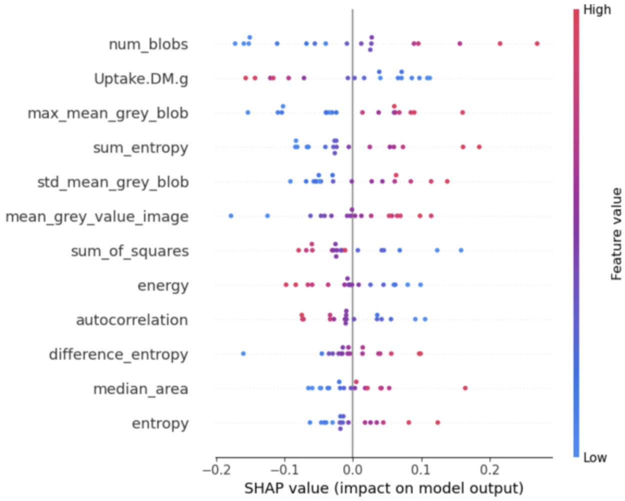


Figure S7: SHAP analysis ranks the top 12 morphological features impacting mean resistance, led by structural complexity and PEDOT:PSS uptake.

To further explore discrete relationships between morphology and resistance, model training and feature importance was performed again with the original morphological feature set (Figure S7). A model was resolved with a coefficient-of-determination of $R^2$=0.43, from which below shows the results of feature importance ranking via SHAP analysis displaying the top 12 features based on magnitude of impact on mean resistance.

The top two ranked features for predicting mean resistance are the number of discrete blobs detected in the image and the absolute mass of PEDOT:PSS taken up by the specimen. `'num_blobs'` is the number of detected blobs, representing the density of distinct features or objects in the image and can indicate fragmentation, clustering, or structural complexity. This would suggest that the combination of complexity with uptake mass of PEDOT:PSS is impactful to resistance, with high resistance associated with increased complexity and reduced uptake.

Additional features (in descending importance) are:

1. Maximum mean grey value, or the within-blob maximum tissue density.
2. Textural complexity `'sum_entropy'`, with high textural complexity associated with increased resistance.
3. Within-blob mean grey value variance, indicating high tissue density variance within blobs as associated with increased resistance.
4. The global mean grey value of the specimen.
5. Image texture as sum of squares, where low values would indicate more homogeneous or smooth textures.
6. Texture uniformity as energy, where low values would indicate complex textures.

7. Autocorrelation indicating the degree of repetitive patterns or spatial similarity, with low values indicating more random and irregular textures.
8. Texture as difference entropy, with high values indicating greater texture complexity.
9. The median size of detected blobs, indicating greater resistance with larger blob sizes.
10. Texture complexity as entropy, where high values are associated with increased texture complexity.

In total, the feature importance narrative suggests that morphological complexity is highly consequential to ultimate resistance in combination with the absolute mass of PEDOT:PSS taken up by the specimen and density of the specimen. In this case it is suggested that as morphological complexity increases resistance increases independent of the absolute amount of PEDOT:PSS infused. Given this, it may be suggested that a range of resistance or electrical responsiveness may be accessed as a function of morphology given the wide range of morphological complexity and variation addressable in aerial mycelium.

##### 1.4 Classification of Local Electrical Properties of Mycelium Chip Specimens as a Function of Global and Local Morphological Qualities

To identify meaningfully distinct mycelium morphologies for characterization, the source aerial mycelium sample population was processed with the morphological featurization pipeline and discretized using Principal Component Analysis (PCA) and k-means clustering [11, 12]. K-means clustering was employed to identify patterns in the extracted image features by partitioning the data into K distinct clusters based on similarity. Prior to clustering, features were standardized using z-score normalization to ensure equal weighting. The optimal number of clusters was selected via the elbow method, which evaluates the within-cluster sum of squares (WCSS) across increasing values of K. PCA was applied for dimensionality reduction and visualization of the clustering results. Clusters were selected by first identifying the pair of clusters with the greatest Euclidean distance between their centroids, representing the most dissimilar pair. A third cluster was then chosen by maximizing the combined distance from both of the initially selected clusters, ensuring maximal overall diversity among the three selected clusters. The below plot shows morphologically distinct cluster groups within the k-dimensional space (PC1 and PC2), where specimens from clusters 0, 2, and 3 were selected for subsequent characterization in order to maximize morphological variety (Figure S8).

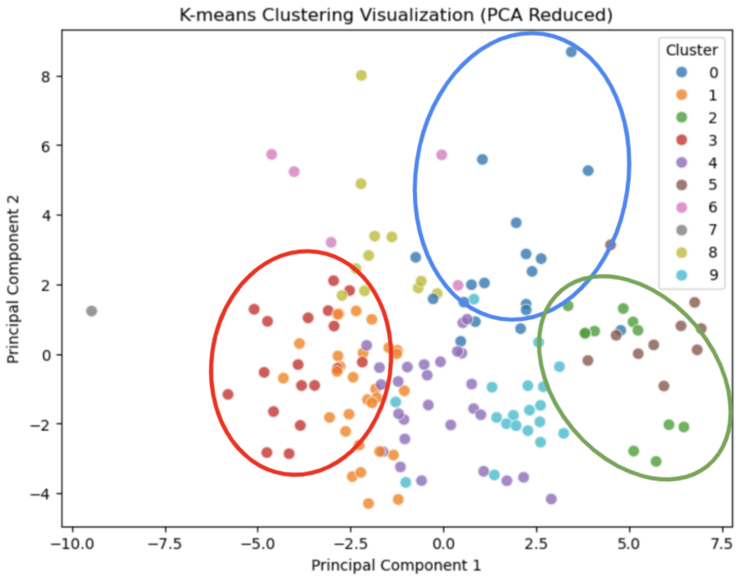


Figure S8: PCA and k-means clustering of morphological features identify three maximally distinct mycelium specimen groups selected for downstream electrical characterization.

We were able to cluster the original mycelium mat sample sets discretized into morphologically related cluster groups 0, 2, and 3 per the above k-means cluster analysis (Figure S9).

Finally, mycelium chip specimens were prepared from one representative mycelium mat specimen from each of the selected morphological groups (Figure S10). Specimens were prepared in acrylic templates such that local electrical

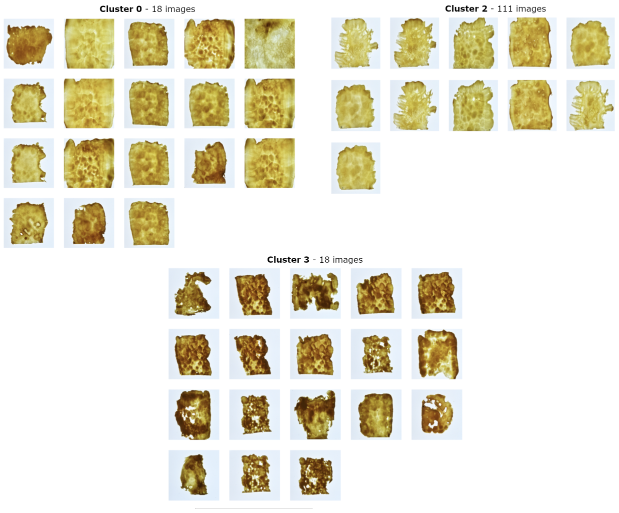


Figure S9: Mycelium samples clustered into groups 0, 2, and 3 based on morphological similarity.

and morphological properties could be evaluated at up to 10 discrete positions. The below image shows the prepared 100x50x1mm specimens from each of classes 0, 2, and 3 before and after PEDOT:PSS infusion.

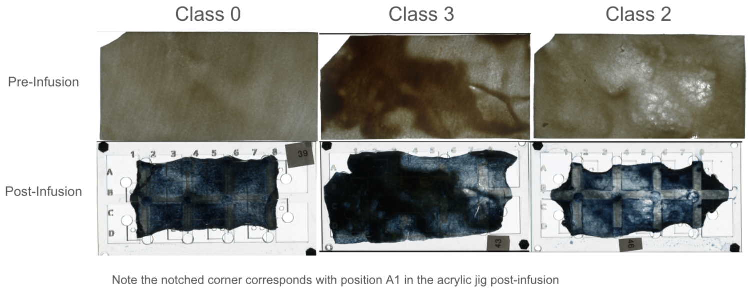


Figure S10: Representative mycelium chip specimens from clusters 0, 2, and 3 before and after PEDOT:PSS infusion, prepared for localized electrical and morphological analysis.

Local electrical properties were characterized according to the sample positions defined by the acrylic template. Mycelium chip specimens were imaged before and after PEDOT:PSS infusion, images cropped to their respective final dimensions, and pre- and post-infusion images aligned using a rigid body alignment in FIJI [13, 14]. Regions of interest (ROI) were defined based on the acrylic sampling template in the post-infusion image, then each ROI measured from the pre-infusion specimen in order to correlate local post-infusion electrical properties to pre-infusion tissue morphology. For each ROI the grey value distribution and summary statistics were measured in FIJI. Histogram-based image features were extracted from the greyscale intensity distributions by computing statistical (mean, variance, skewness, kurtosis), shape (entropy, energy, contrast, homogeneity), and cumulative (interquartile range) descriptors. Additionally, whole chip specimen morphological featurization was performed as previously and dimension reduced via PCA to provide global morphological features. Resistance and conductivity data from electrical characterization were aggregated and the IV curves featurized by fitting linear regression models for each ROI, extracting parameters including slope, intercept, $R^2$, p-value, and residual statistics. Finally, electrical target distributions were evaluated and appropriate transformations (boxcox or log) applied to manage outliers and skewness prior to model training. This resulted in a data set from which local electrical properties after PEDOT:PSS infusion could be evaluated as a combined function of global and local mycelium morphology prior to infusion. Subsequent analysis evaluated the potential for explaining post-infusion local electrical properties based on pre-infusion morphology.

Figure S11 illustrates ROI alignment to pre- and post-PEDOT:PSS infused specimens (morphological class 0), as well as characterized current (2V) and I–V non-linearity (sum of squared residuals).

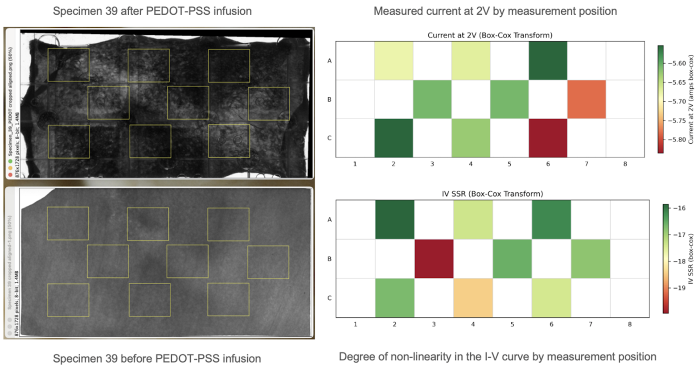


Figure S11: ROI alignment on pre- and post-infusion specimens (class 0), with mapped current (2V) and I–V non-linearity illustrating spatial variation in electrical response.

PCA was performed on the target space (combination of all conductance, resistance, and I–V features) to determine the degree of co-variability. The results suggest that all measured electrical features are highly correlated with one another, collapsing the whole electrical response space to a single response (‘PC1’, explaining >90% of total electrical variance) (Figure S12).

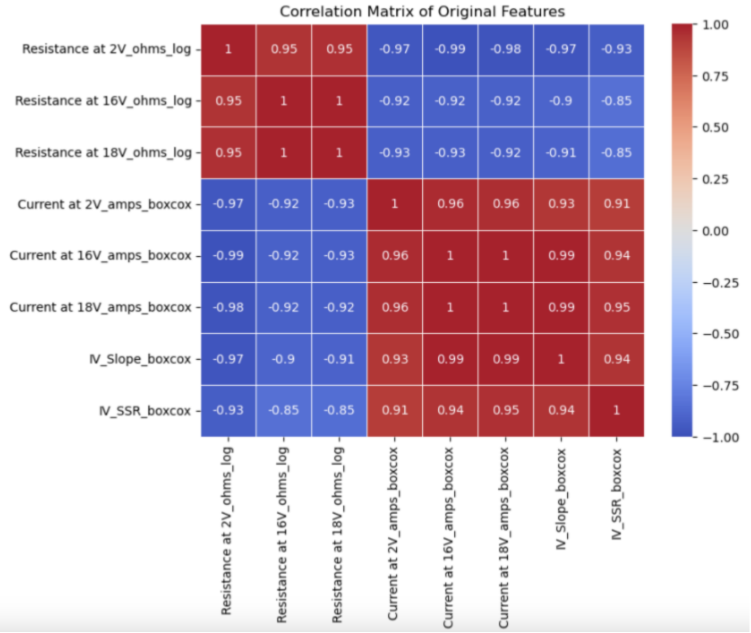


Figure S12: PCA of electrical features shows strong covariability, with PC1 capturing over 90% of total variance across conductance, resistance, and I–V metrics.

Mutual information analysis was performed to quantify the dependency between each of 17 features and multiple target variables using the ‘`mutual_info_regression`’ function from Scikit-learn[15, 16, 17]. For each target, mutual information scores were computed, stored, and visualized to highlight the most informative features. The results help identify which features share the most information with each target, where higher scores suggest a stronger relationship. An aggregated average score across all targets was used to assess overall feature relevance. The results suggest that features are not noise-dominated and that the feature space holds predictive value for the electrical property target variable (Figure S13). Suggested feature importance in descending order is (1) global morphology (density and complexity), (2) local tissue density, (3) local tissue complexity/uniformity and eccentricity.

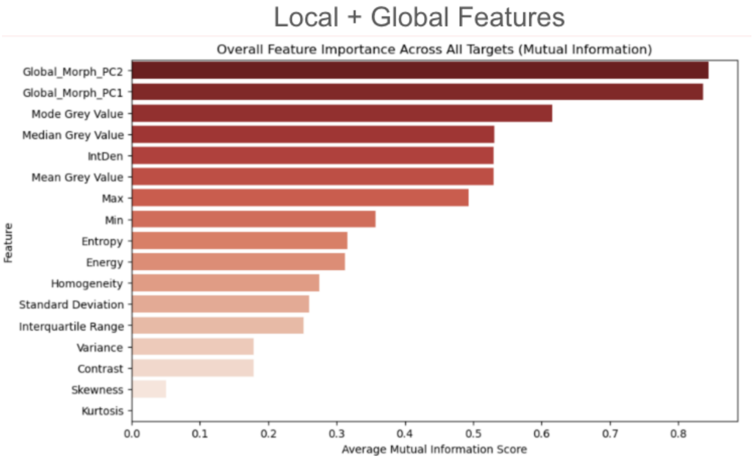


Figure S13: Mutual information analysis identifies global morphology, local density, and texture complexity as top predictors of electrical properties.

A regression neural network was trained to predict electrical property PC1 as a combined function of both global and local morphological features. A fully connected feedforward neural network was trained using leave-one-out cross-validation (LOOCV) to predict the electrical property (PC1) target from the feature set. The model architecture consisted of two hidden layers with 128 and 64 ReLU-activated units, respectively, incorporating L2 regularization and dropout to mitigate overfitting. Model performance was evaluated by aggregating prediction error and loss across all LOOCV folds, with mean squared error as the loss function and the Adam optimizer used for training.

A coefficient-of-determination of 0.74 was achieved, suggesting strong predictive utility for endpoint electrical properties as a function of original tissue morphology, allowing for classification of probable local electrical properties across an arbitrary morphological surface prior to physical infusion with PEDOT:PSS.

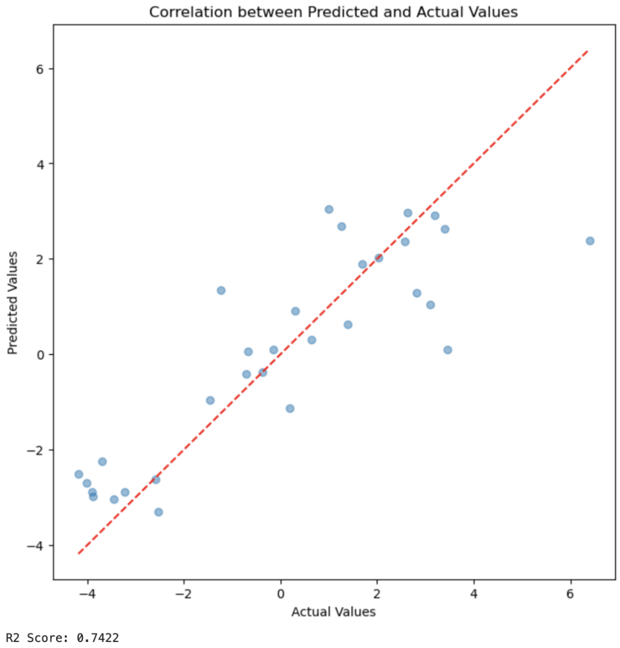


Figure S14: Mutual information analysis identifies global morphology, local density, and texture complexity as top predictors of electrical properties.

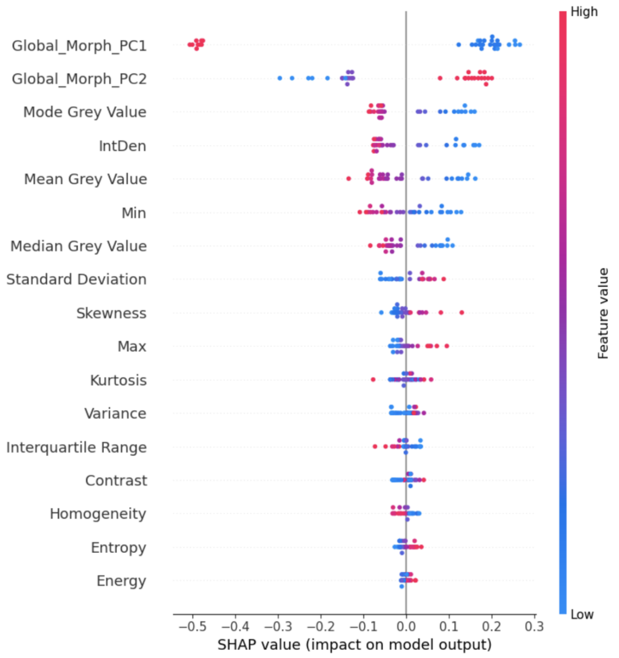


Figure S15: SHAP analysis ranking feature importance for electrical property response PC1 values.

The results suggest that electrical performance of PEDOT:PSS infused mycelium chips have a significant relationship with morphological variation within the feedstock mycelium mat prior to PEDOT:PSS infusion. Further, the distribution of local electrical properties over a given mycelium mat area may be classified prior to infusion with PEDOT:PSS. The ability to pre-classify the distribution of electrical properties across a mycelium sheet prior to PEDOT:PSS infusion offers a powerful means of functionally screening and selecting material regions with desirable electrical behavior for reservoir computing depending on use case and application. By predicting properties such as resistance and I–V non-linearity based on native morphological features, this approach enables targeted use or exclusion of specific regions, optimizing the functional capacity and consistency of the reservoir. This pre-infusion classification may reduce empirical trial-and-error in reservoir training and support rational design of mycelium-based computational substrates through targeted morphological engineering during growth.

##### 1.5 Detailed Statistical Analysis of Temporal Dynamics

We assessed temporal dynamics on a single device using random ($K = 9$ technical runs), sinusoidal ($K = 3$ runs), and step ($K = 2$ runs) inputs, chosen to probe complementary memory regimes.

###### Random Input Analysis

For random inputs, a linear readout using only the current input served as a baseline. Adding short input history (input($t \ldots t - 10$)) improved next-step state prediction within-device: mean $\Delta R^2 = 0.107$, 95% CI 0.048–0.166; paired two-tailed $t(8) = 3.33$, $p = 0.010$, Cohen's $d = 1.11$. This indicates shallow but reliable carryover on sub-second to $\sim$1 s scales.

###### Correlation Structure:

- State autocorrelation: median peak ACF $= 0.342$ at 2.5 s (positive lags only)
- Input–state coupling: median CCF $= 0.721$ at 0 s (non-negative lags)
- Interpretation: Strong feedthrough with modest persistence

###### Sinusoidal Drive Analysis

Under sinusoidal drive, the state exhibited phase-locked memory aligned to the waveform:

- Median peak ACF $= 0.497$ at 1.0 s
- Median peak CCF $= 0.362$ at 1.0 s (lags in seconds from median sampling interval)

###### Predictive Performance:

- Current input only: mean $R^2 = 0.184$, 95% CI 0.000–0.470
- With input history: mean $R^2 = 0.230$, 95% CI 0.000–0.475
- Improvement: mean $\Delta R^2 = 0.045$, 95% CI 0.000–0.131
- Statistical test: $t(2) = 1.06$, $p = 0.40$ (not significant)

**Run-by-Run Variability:** Of three sine runs on the same device, two showed clear phase-locked input–state coupling (CCF peaks 0.36–0.69 at 0–1 s); one run was non-responsive (max CCF $\approx 0$, $R^2 \approx 0$). All runs were retained in primary analysis (Table 1).

###### Step Input Analysis

Step inputs revealed multi-timescale dynamics:

- Median peak ACF $= 0.982$ at 3.15 s
- Median peak CCF $= 0.472$ at 11.1 s

###### Standard Step Response Metrics:

- $\tau \approx 0.15$ s (first crossing of 63.2% of effective step)
- 10–90% rise $\approx 4.35$ s
- Overshoot $\approx 143\%$ relative to late-window mean
- No $\pm 10\%$ settling within recorded windows

**Representative Behavior:** The system crossed 63.2% within a single sampling interval yet continued slow drift and overshoot thereafter, indicating a rapid channel superimposed on slower relaxation that prevents classical first-order settling over available duration.

**Methodological Definitions**

- Rise time: 10–90% of effective step amplitude
- $\tau$: First crossing of 63.2%
- “Steady”: Mean over final 20% of each trace
- Settling: Earliest time state enters and remains within $\pm10\%$ (and $\pm5\%$) of steady value
- Correlations: Pearson coefficients
- ACF peaks: Reported at positive lags
- Input–state CCF peaks: Reported over non-negative lags with lags converted to seconds
